# Efficient CRISPR/Cas-mediated homologous recombination in the model diatom *Thalassiosira pseudonana*

**DOI:** 10.1101/215582

**Authors:** Nigel Belshaw, Irina Grouneva, Lior Aram, Assaf Gal, Amanda Hopes, Thomas Mock

## Abstract

CRISPR/Cas enables targeted genome editing in many different plant and algal species including the model diatom *Thalassiosira pseudonana*. However, efficient gene targeting by homologous recombination (HR) to date is only reported for photosynthetic organisms in their haploid life-cycle phase and there are no examples of efficient nuclease-meditated HR in any photosynthetic organism. Here, a CRISPR/Cas construct, assembled using Golden Gate cloning, enabled highly efficient HR for the first time in a diploid photosynthetic organism. HR was induced in *T. pseudonana* by means of sequence specific CRISPR/Cas, paired with a donor matrix, generating substitution of the silacidin gene by a resistance cassette (FCP:*NAT*). Approximately 85% of NAT resistant *T. pseudonana* colonies screened positive for HR using a nested PCR approach and confirmed by sequencing of the PCR products. The knockout of the silacidin gene in *T. pseudonana* caused a significant increase in cell size, confirming the role of this gene for cell-size regulation in centric diatoms. Highly efficient gene targeting by HR makes *T. pseudonana* as genetically tractable as *Nannochloropsis* and *Physcomitrella*, hence rapidly advancing functional diatom biology, bionanotechnology and any biotechnological application targeted on harnessing the metabolic potential of diatoms.

## Introduction

Diatoms represent a highly successful group of unicellular phytoplankton responsible for an estimated 20% of global primary production (Field et al. 1998; Falkowski and Raven 2007). Due to their silicified cell walls (frustules) diatoms also dominate the ocean’s biogenic silicon cycle (Treguer and De La Rocha 2013). Many of the traits underpinning their ecological success are the reason why these eukaryotic microbes are desirable for algal biotechnology and fuel production (Bozarth, Maier, and Zauner 2009; Levitan et al. 2014; Wang and Seibert 2017). Their most characteristic traits include fast growth, biomineralisation and complex metabolism derived from a) successive endosymbiotic events (Bhattacharya et al. 2007; Frommolt et al. 2008; Moustafa et al. 2009; Keeling 2010), b) genome hybridization (Tanaka et al. 2015) and c) haplotype divergence (Paajanen et al. 2017; Mock et al. 2017). Interestingly, the fastest specific cell division rate for any autotroph (0.54 divisions per hour) was measured in a centric marine diatom: *Chaetoceros salsugineum* (Ichimi et al. 2012). The processes that govern this remarkable ability are largely unknown but are probably shared by many diatoms as they generally outcompete other algal groups when growth conditions become favourable. Their metabolism was shaped by different endosymbiotic events (primary and secondary endosymbiosis) that took place at least 1 billion years ago and contributed various different enzymes, isoforms and entire pathways such as the urea pathway (Armbrust et al. 2004; Allen et al. 2011). These ‘mix and match’ genomes are considered to have contributed to the plasticity of their genomes as well as phenomes and therefore their trait performance landscapes (Messina et al. 2011). More recent evolutionary processes that shaped diatom genomes include genome hybridisations (Tanaka et al. 2015) and haplotype divergence (Paajanen et al. 2017; Mock et al. 2017). The latter is considered to be the consequence of adaptive evolution leading to environment-dependent Differential Allelic Expression (DAE) (Paajanen et al. 2017; Mock et al. 2017). The correlation between diversifying selection and allelic differentiation suggests a potential role of the divergent alleles for adaptation to environmental fluctuations in aquatic environments.

To harness the unique biology of diatoms for advancing fundamental research and algal biotechnology, we need to improve our knowledge of molecular factors controlling their evolution, adaptation and metabolism. Genome-enabled studies have revealed genes, regulatory elements and epigenetic factors potentially controlling the expression of diverse phenotypes. It is only recently, however, that genome editing tools such as CRISPR/Cas (Sander and Joung 2014; Doudna and Charpentier 2014; Lander 2016) and TALENs (Gaj, Gersbach, and Barbas 2013) have become available, including in diatoms (Daboussi et al. 2014; Hopes et al. 2016; Nymark et al. 2016; Weyman et al. 2015; Serif et al. 2017), enabling us to improve our understanding of individual genetic factors for the evolution and biology of diatoms. The introduction of gene-targeting by high frequency HR allows precise genetic changes via knock-in of a particular sequence at a specific locus, rather than just gene knock-out. Thus, a proven ability of gene targeting via HR in diatoms will enable improvements in gene and protein engineering, which will be instrumental to advance our understanding of fundamental diatom biology and to facilitate diatom domestication.

Gene targeting via HR has been demonstrated in a number of photosynthetic eukaryotes to date including *Cyanidioschizon merolae* (Minoda et al. 2004), *Ostreococcus tauri* (Lozano et al. 2014), *Chlamydomonas reinhardtii* (Greiner et al. 2015), *Nannochloropsis sp.* (Kilian et al. 2011), *Phaeodactylumm tricornutum* (Daboussi et al. 2014; Weyman et al. 2015)*, Physcomitrella patens* (Kamisugi et al. 2006; Kamisugi, Cuming, and Cove 2005; Kamisugi, Whitaker, and Cuming 2016), *Arabidopsis* (Schiml, Fauser, and Puchta 2014), *Oryza* (Begemann et al. 2017) and *Nicotiana* (Li et al. 2013). Most of these, however, are either haploid and/or have low rates of HR. Highly efficient HR can be seen in two of these species: the moss *Physcomitrella patens* and the eustigmantophyte *Nannochloropsis sp.*. *Nannochloropsis sp*. is haploid, and the dominant phase of the life cycle in *P. patens* is also haploid, which makes backcrossing to establish homozygous transgenic lines obsolete. Targeted and efficient HR in both species was achieved without introducing nucleases to produce DNA double-strand breaks. For example, in *Nannochlorpsis sp*., Kilian et al. (2011) used 1 kb flanking sequences to target genes for HR-induced gene knock-out. For all the other photosynthetic eukaryotes, rates of HR are rather low either with or without the help of nucleases. There are two reports of TALEN-mediated gene editing including HR in the marine diatom *P. tricornutum* (Daboussi et al. 2014; Weyman et al. 2015). Both studies tested the occurrence of HR by co-transformation of the TALEN-carrying plasmid and a plasmid with a donor template and reported an efficiency of up to 27% (Daboussi et al. 2014; Weyman et al. 2015).

The present study, however, is the first report of highly efficient HR in any diploid photosynthetic organism mediated by CRISPR/Cas. The aim of this work was to show that HR can be induced in a diatom by means of the sequence-specific nuclease Cas9, paired with a donor matrix (exogenous DNA templates with homology to the targeted locus), generating a complete knock-out of a target gene in *T. pseudonana*. In this study, the silacidin gene was substituted by a resistance cassette (FCP:*NAT*). By extension, this method could also enable targeted gene insertion/substitution in *T. pseudonana* and possibly other diatoms as the versatile Golden Gate cloning system was used (Weber et al. 2011; Belhaj et al. 2013) to make the vector constructs. High frequency gene targeting by HR will considerably expand the genetic tool set for diatoms and has already made *T. pseudonana* as genetically tractable as *Nannochloropsis* and *Physcomitrella*, hence rapidly advancing diatom functional genomics and biotechnology.

## Material and Methods

### Strains and growth conditions

*Thalassiosira pseudonana* (CCMP 1335) was grown in half salinity (½ SOW: 16 g l^−1^) Aquil sea water medium (Price et al. 1989) supplemented with silica under constant illumination (ca. 100 µmol photons m^−2^ s^−1^) at 20°C.

### Plasmid design and assembly

Two plasmids were used to introduce all components necessary for HR (Fig. 1). 1) The first plasmid contained Cas9 and one single guide RNA (sgRNA) designed, due to repetitive nature of the silacidin sequence, to induce two cuts within the silacidin gene (pAGM4723_Tp:FCP_P:Cas9:T_TpU6P:sgRNA). 2) The second plasmid contained a cassette including the nourseothricin resistance gene *NAT*, flanked by two regions homologous to non-coding 3’ and 5’ ends (629 bp and 868 bp) of the silacidin gene to serve as a template (donor matrix) for HR. The second plasmid was assembled from PCR products equipped with BpiI restriction sites and overhangs for directional cloning (pAGM4723_5’Sil_TpFCP:NAT_3’Sil) to allow cloning by the Golden Gate method. Cloning was carried out according to Belhaj et al. (2013) and Weber et al. (2011) in a single-step restriction/ligation reaction. Forty fmol of each component was combined in a 20 μl reaction with 10 units of BsaI (L1 assembly) or BpiI (L2 assembly) and 10 units of HC T4 DNA ligase in 1 × ligation buffer. The reaction was incubated at 37 °C for 5 h, 50 °C for 5 min and 80 °C for 10 min. Five μl of the reaction was transformed into 50 μl of NEB 5-alpha chemically competent *E. coli*.

**Figure 1.**
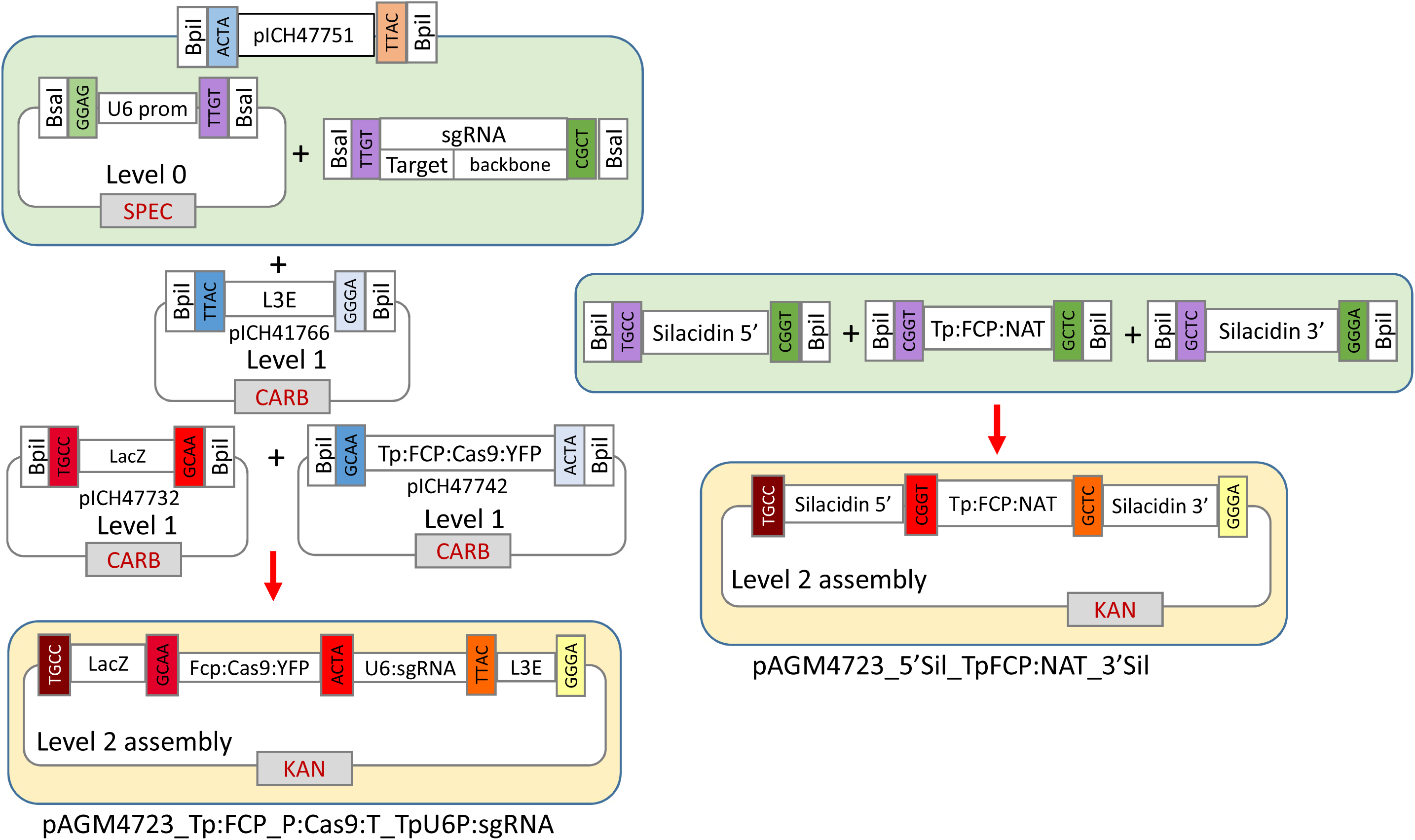
Overview of modules and overhangs used in Golden Gate assembly of level 2 transformation vectors pAGM4723 Tp:FCP_P:Cas9:T_TpU6P:sgRNA and 5’Sil_TpFCP:NAT_3’Sil. Tp:FCP: module flanked by endogenous *T. pseudonana* FCP gene promoter and terminator sequences. Plasmids carry antibiotic resistance selection genes against: spectinomycin (SPEC), carbenicillin (CARB) and kanamycin (KAN).

### Plasmid 1 assembly

The sgRNA was amplified with primers 1 (introducing the specific 20 nt region at the 5’ end) and 2 (Table 1) using plasmid pICH86966_AtU6p_sgRNA_NbPDS (containing the sgRNA backbone) as a template (Table S1). PCR was carried out with Phusion DNA polymerase with 56°C annealing temperature and 1 min elongation step. The purified PCR product was then assembled together with the L0 vector containing the Tp:U6 promoter sequence (Hopes et al. 2016) into the L1 vector pICH47751. L1 modules pICH47732 (lacZ), pICH47742 (CAS9), pICH47751:TpU6p:Sil_sgRNA, pICH41766 (L3E) were then assembled together in the L2 backbone pAGM4723. For details on sequences, restriction sites and selection markers see Addgene data base, Hopes et al. 2017 and table S1.

**Table 1.**
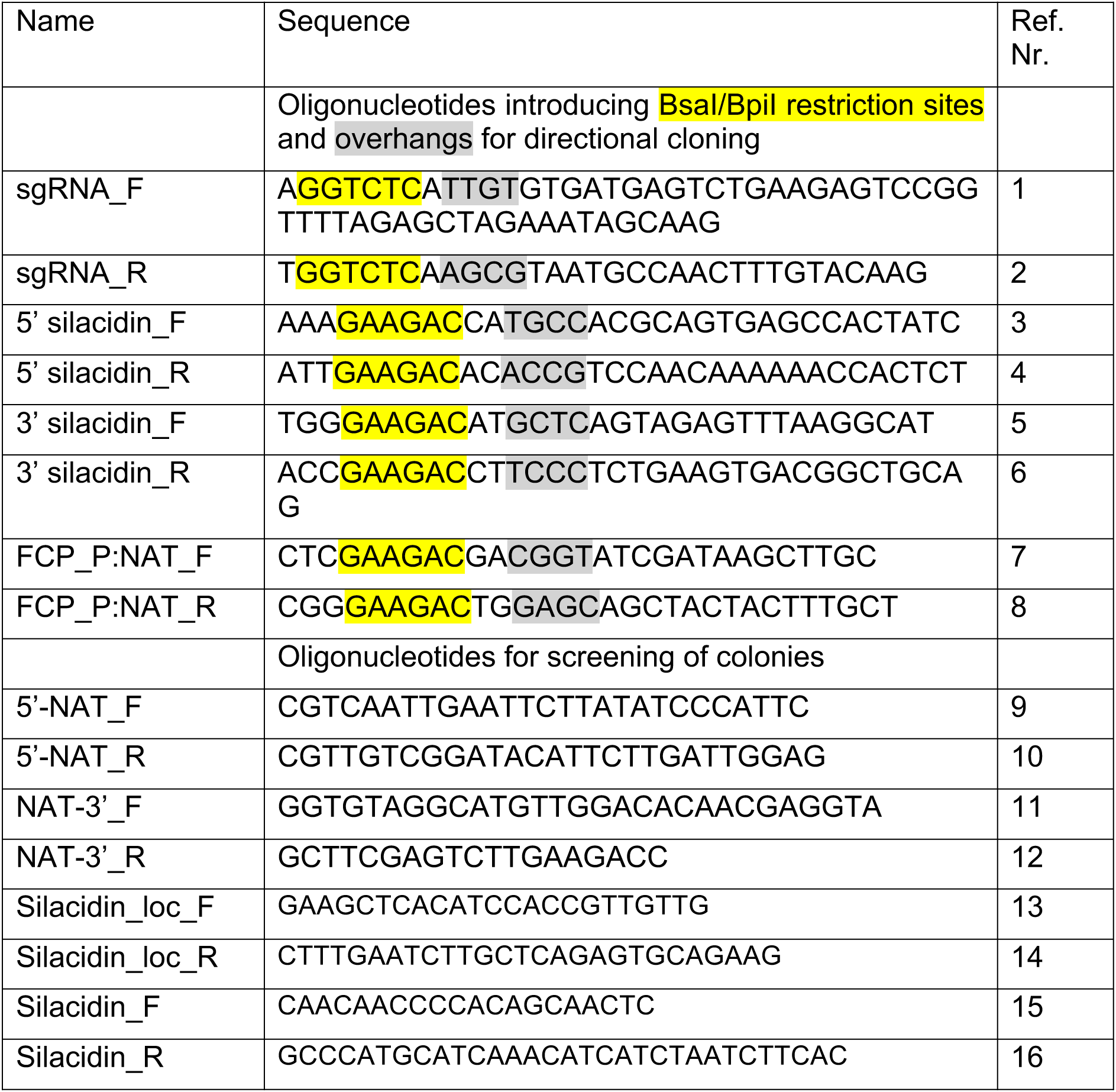
Oligonucleotides used for cloning and screening

### Plasmid 2 assembly

5’ and 3’ silacidin flanking, non-coding sequences were amplified from genomic DNA with primer pairs 3/4 and 5/6 introducing overhangs for directional cloning and BpiI restriction sites (Table 1). The NAT resistance gene cassette with *T. pseudonana*-specific promoter and terminator was amplified with primers 7/8 with vector pICH47732:FCP:NAT as template (Hopes et al. 2016). Phusion DNA polymerase was used and the annealing temperature was 56°C in all cases. After purification, PCR products were assembled into a pAGM4723 backbone using the Golden gate method described above.

### sgRNA selection and design

The silacidin gene is composed of extensive repeats. The sgRNA used in the present work (GTGATGAGTCTGAAGAGTCCGGTTTTAGAGCTAGAAATAGCAAGTTAAAATAAG GCTAGTCCGTTATCAACTTGAAAAAGTGGCACCGAGTCGGTGC*TTTTTT*, where the underlined sequence indicates the 20nt target), was designed to induce two cuts within the coding region of the gene. It has to be noted, however, that two variants of silacidin (allele variation) are present in the database. Only one of them predicts two identical target sequences matching the seed sequence of the selected sgRNA. Single guide RNAs were designed according to Hopes et al. (2016) and Hopes et al. (2017).

### *Transformation of* T. pseudonana

Biolistic particle delivery transformation was carried out according to Poulsen et al. (2006) and Hopes et al. 2017).

For each shot, 5 × 10^7^ cells from the exponential growth phase were collected onto a filter (47mm diameter 1.2um Whatman Isopore) and placed on top of a 1.5% agar, ½ SOW Aquil medium petri dish. One µg of each plasmid (Tp:Cas9:sgRNA and Sil_5’:NAT:3’) was used for coating 3 mg of M10 (0.7 µm diameter) tungsten particles (BioRad, Hercules, CA) per shot. A 7 cm flight distance and 1350 psi rupture discs were used. After transformation, each filter was immediately placed into 25 ml of fresh Aquil medium and cells were left to recover under standard growth conditions for 24 h. The following day, 5 × 10^6^ cells were spread onto selective 0.8% ½ SOW Aquil agar plates with 100 µg ml^−1^ nourseothricin. An aliquot of 10^6^ cells ml^−1^ was supplemented with nourseothricin and grown as a liquid culture. Colonies appeared on selective plates after 8-12 days.

### Screening of colonies

Colonies were re-streaked onto fresh selective plates and grown in liquid medium for DNA isolation. DNA was then used as a template for PCR analysis using MyTaq DNA polymerase (Bioline). A nested PCR approach was adopted using primers detailed in table 1. Initially, a primer pair (13/14) targeting the silacidin locus outside the 5’ and 3’ regions used for inducing HR was used to amplify the silacidin locus irrespective of whether HR had occurred. The products of this PCR were used as templates in two PCRs with primers nested to those used in the primary PCR but still 5’ or 3’ to the silacidin flanking regions used in the Sil_5’:NAT:3’ plasmid (9 and 12), together with primers specific for the *NAT* gene cassette (10 and 11, Fig. 2). Specific amplicons from these PCRs signified HR-mediated replacement of silacidin. In addition, a fragment of the silacidin gene including both putative Cas9 cut sites was amplified in a further nested PCR with primers 15 and 16. Substitution by HR resulted in no amplicon, while Cas9:sgRNA-induced deletions resulted in smaller products.

**Figure 2.**
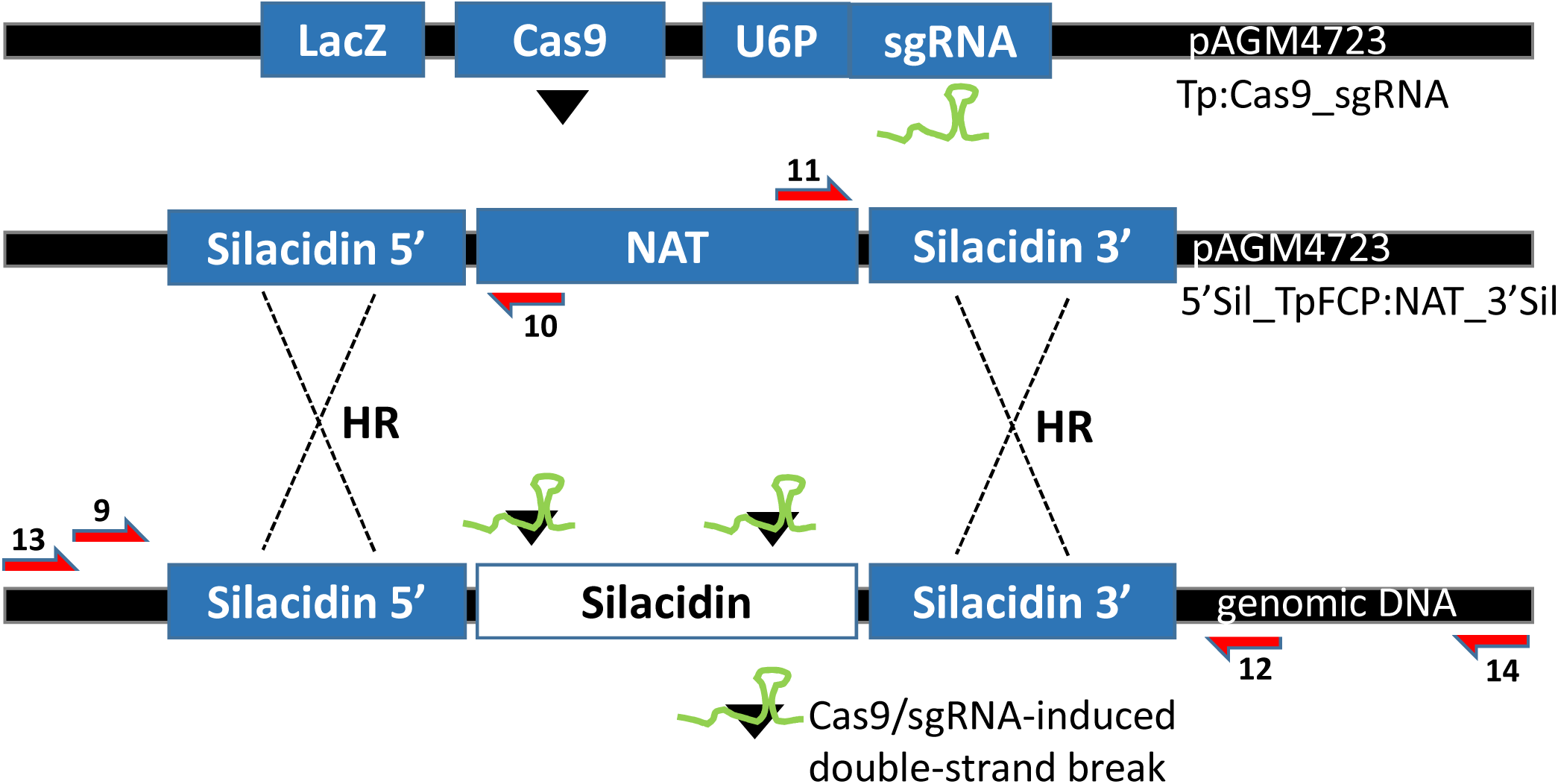
Overview of HR mechanism between donor matrix and genomic silacidin locus in *T. pseudonana*. Vector Tp:Cas9_sgRNA expressed Cas9 nuclease and sgRNA targeting silacidin. Vector 5’Sil_TpFCP:NAT:3’Sil carried the exogenous homology matrix containing *NAT* resistance cassette and non-coding 5’/3’ flanking regions. Red arrows depict primers used in screening strategy by PCR, numbers correspond to table 1.

### Phenotyping based on cell size estimation

In accordance with expectations from earlier experiments on silacidin knock-down mutants that showed larger cell size/valve diameter in transformant lines (Kirkham et al. 2017), we used three different approaches to compare cell sizes of selected mutant lines to WT. All measurements were done on cells during mid-exponential growth phase. i) Culture aliquots were measured with a Z2 Coulter counter (Beckman Coulter). The cell size was determined as the maximum value of the Gaussian distribution of cell sizes in each measurement. ii) Forward scattering of light by single cells was measured with an Eclipse iCyt flow 531 cytometer (Sony Biotechnology Inc., Champaign, IL, USA), equipped with 405 nm and 488 nm solid state air cooled lasers, both with 25 mW on the flow cell and with standard 533 filter set-up. The forward light scattering is proportional to the size of the cell. iii) Direct measurements of the diameter of the valves were done on scanning electron microscope (SEM) images. Sample aliquots were dried on Isopore^TM^ membrane filters (Merck Millipore, Germany), coated with gold, and imaged with a SUPRA55 SEM (Zeiss).

### Genotyping based on PCR amplification of the silacidin gene

ImageJ was used to quantify peak area for each silacidin band, including truncated bands, and each peak of the ladder (100 bp DNA ladder, NEB). The 800 bp band of the ladder was used as a standard. Number of base pairs (bps) was calculated from the mass of the standard (24 ng) and divided by area. This factor was then applied to the area of each silacidin band to calculate the number of bps.

## Results

Eighteen out of 21 *T. pseudonana* transformant clones screened positive for HR (Figs. 3A and B). The entire silacidin locus was amplified with primer pair 13/14 (Table 1) in 21 clones capable of growth on selective medium. These amplicons were used as templates for two further nested PCRs with primer pairs 9/10 and 11/12 (Table 1). The primers were designed to cover the transition from the 3’/5’ end of the silacidin non-coding sequence to the resistance cassette contained within the exogenous HR donor matrix (Fig. 2). The results showed that correct, targeted HR had taken place at both 3’ and 5’ ends between the WT locus and the donor sequence in 85% of cases (Figs. 3A and B). The WT was used as negative control because the amplified combination of up-/downstream non-coding silacidin sequences interrupted by a selective marker does not occur naturally (Fig. 2).

**Figure 3.**
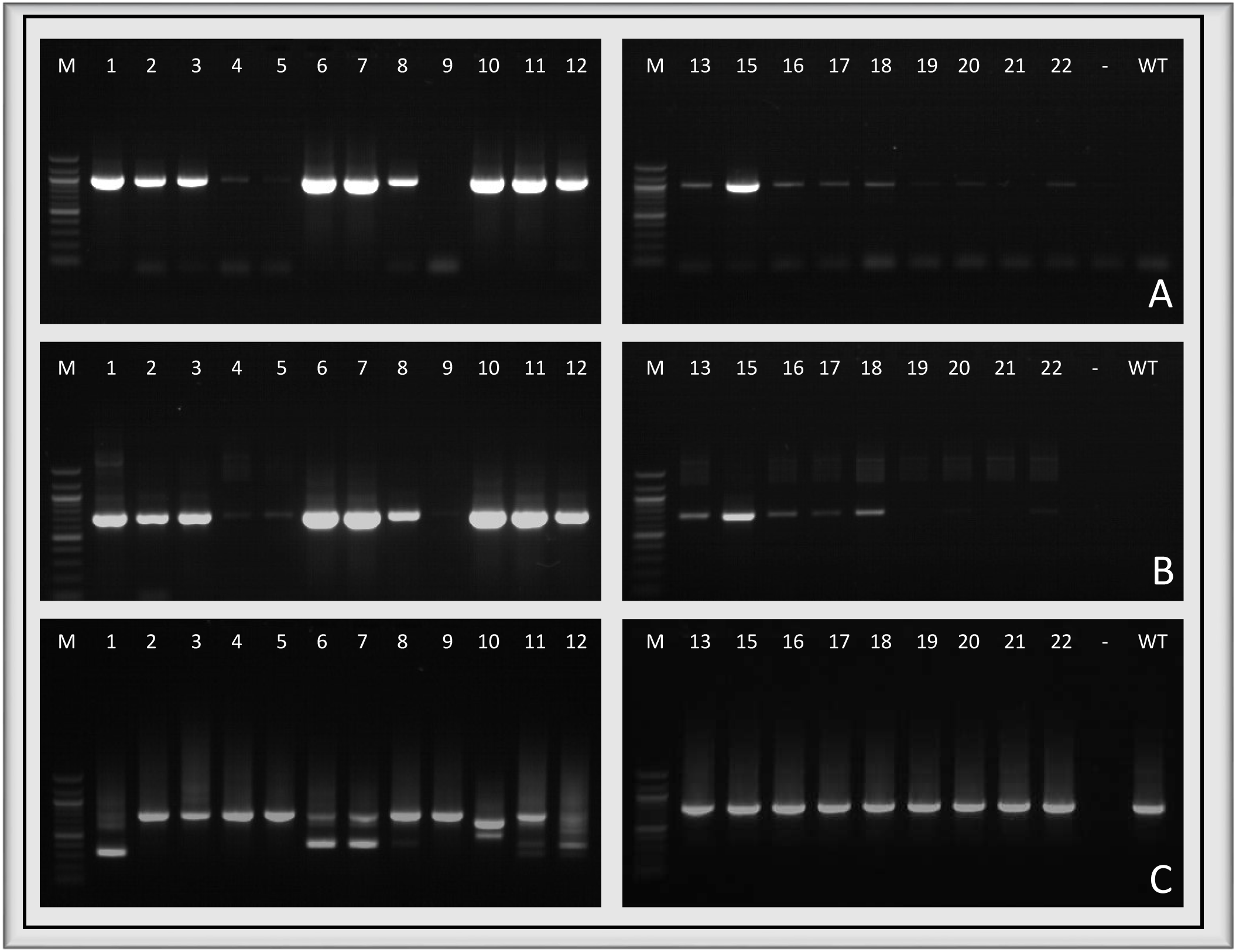
Screening for HR and silacidin deletion by PCR. Primers target genomic region up-/downstream of 5’/3’ flanking regions used for HR construct in combination with *NAT* resistance gene cassette. A: Primer pair 9/10, silacidin locus template. B: Primer pair 11/12, silacidin locus template. C: Silacidin fragment amplicon, primer pair 15/16, genomic DNA template. M: 100bp size ladder. -: no template control. WT: wild type. 1-22: transformant clones.

Screening for the presence of silacidin (PCR with primer pair 15/16) showed that the WT-sized sequence was present in 18 clones and absent in three. In addition, six clones contained shorter bands (Fig. 3C). Sequencing data confirmed that they were all truncated versions of silacidin and not non-specific products (Fig. S1). Shorter amplicons were expected to be the result of Cas9 cutting the sequence and the resulting ends being subsequently repaired by NHEJ. Sequencing data showed that the truncated versions were the result of different deletions within the silacidin sequence (Fig. S1). This was expected because previous studies in diatoms (Nymark et al. 2016; Hopes et al. 2016), showed either small (Hopes et al. 2016) or larger deletions (Nymark et al. 2016) as a result of Cas9 nuclease activity. Furthermore, in plants, other repair outcomes were also shown to be possible (Schiml, Fauser, and Puchta 2014).

Taken together, the above results revealed that in some cases (e.g. clones 6 and 7) three possible outcomes of targeted gene editing were detected per clone (HR, WT silacidin and truncated silacidin). Because *T. pseudonana* is a diploid, only two outcomes are expected to occur per cell, affecting either allele. This suggests that some clones were mosaic.

The presence of a Cas9 amplicon was detected in 20 out of 21 transformants (Fig. 4). The nuclease activity of Cas9 was expected to be a prerequisite for HR induction. The absence of Cas9 in clone 3 (which nevertheless screened positive for HR) indicated the possibility that Cas9 might have been transiently active but subsequently removed from the genome.

**Figure 4.**
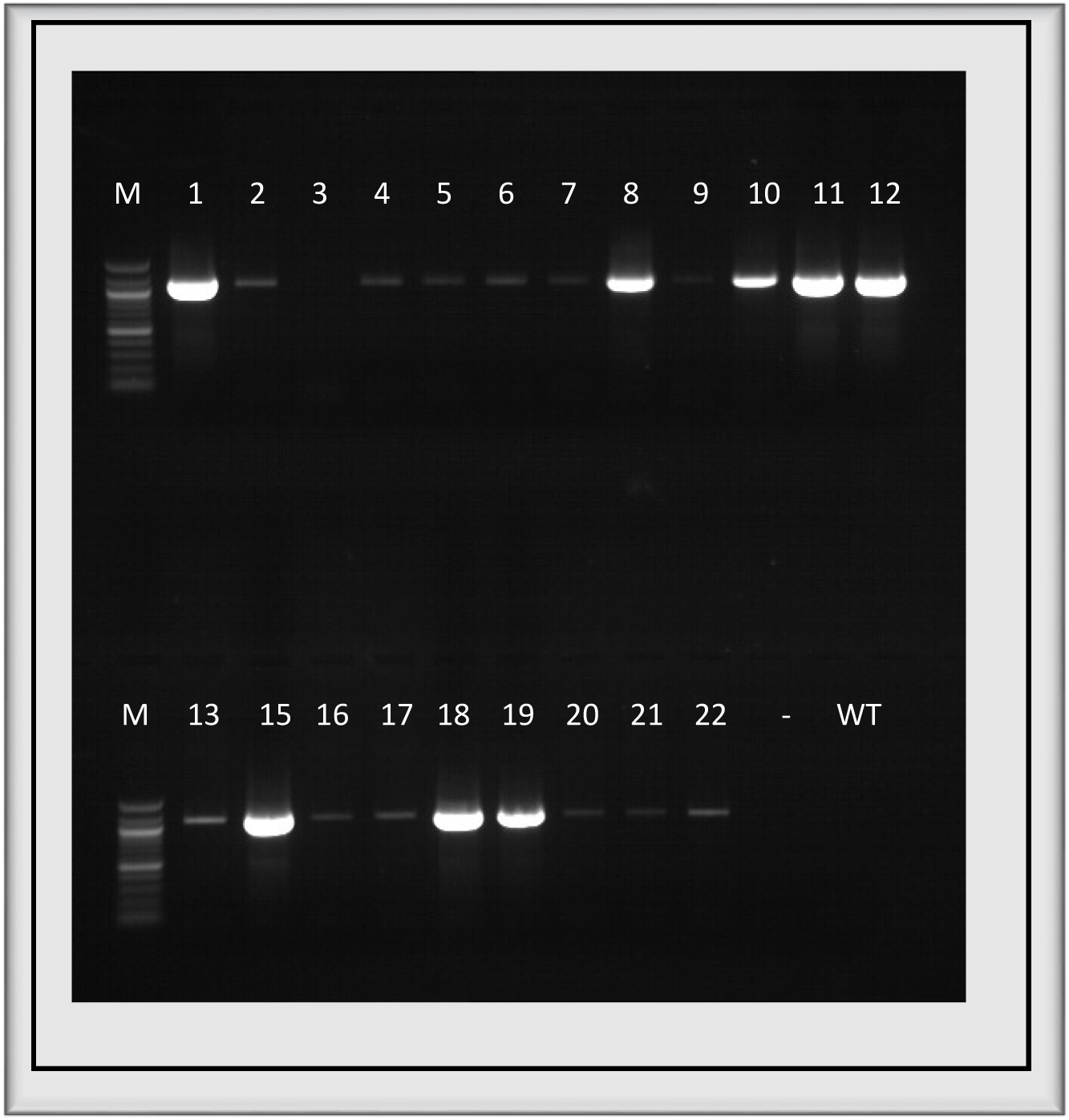
Cas9 fragment PCR. Genomic DNA template. M: 100bp size ladder. -: no template control. WT: wild type. 1-22: transformant HR clones.

Phenotyping of selected clones showed that transformant cells were significantly larger than WT (Table 2). This was most pronounced in clone 12 where a complete deletion of the silacidin gene was identified (Fig. 3C). Quantification of all silacidin PCR products, including truncated versions, from WT and HR clones 1, 6, 7, 10 and 11 gave a significant negative correlation (Fig. 5A, R^2^=0.8, p=0.007) between the quantity of product, given in base pairs, and cell size measured by FACS scatter. For a more detailed analysis of functionality, truncated versions of the silacidin gene were translated into the corresponding amino acid sequence. The resulting polypeptides were aligned with the original WT protein sequence (Fig. 5B). This showed that, if expressed and translated, clones retained a different number of repetitive peptide motifs following the overall amino acid sequence: (SS)SEDSXDSXPSDESEESEDSVSSEDED. This motif was present as a single copy in clone 1, two copies in clone 6 and 7 and four remaining copies in clone 10.

**Table 2.**
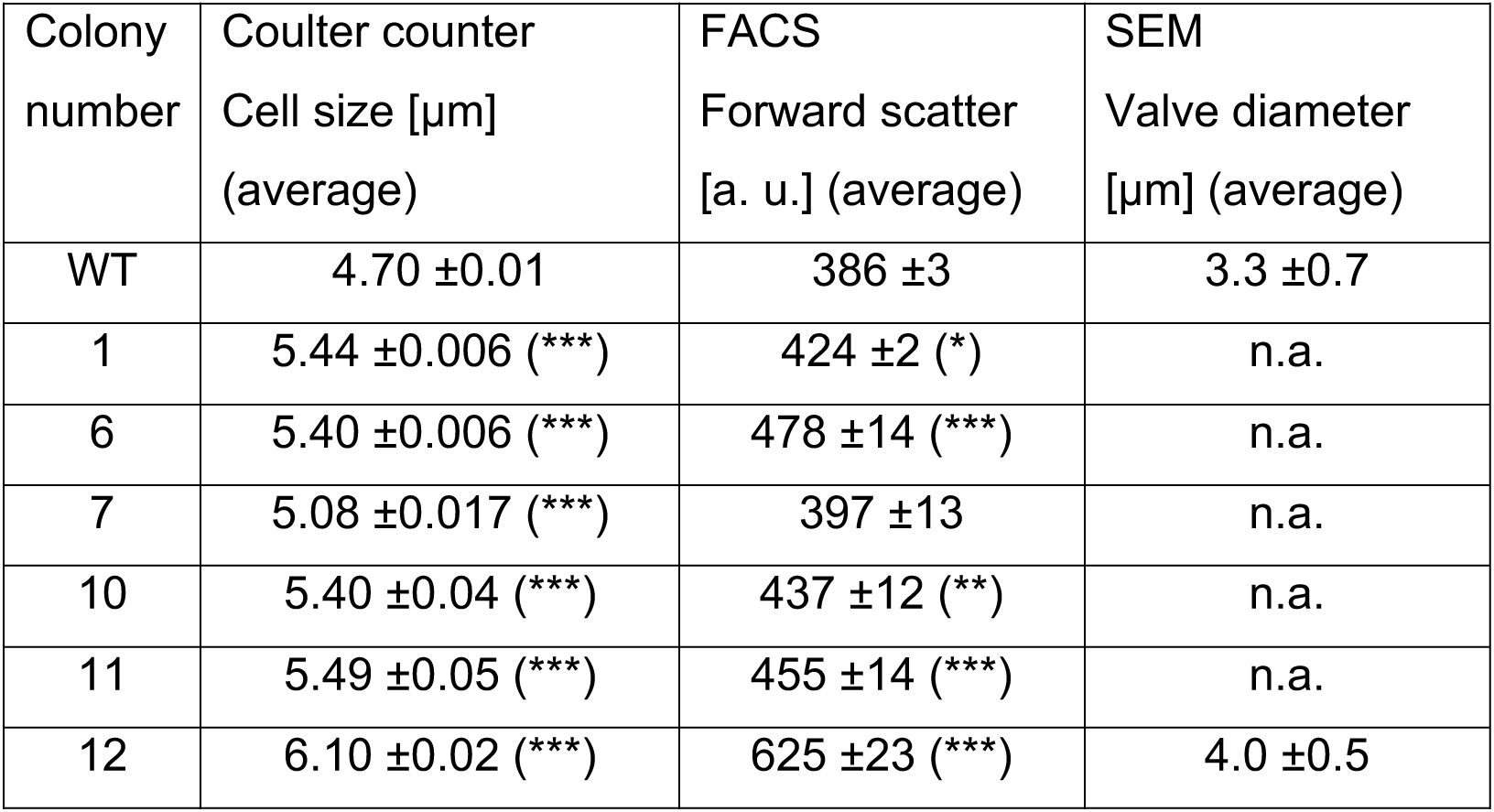
Cell size and valve diameter of *T. pseudonana* WT and transformant lines (1, 6, 7, 10, 11, 12). Average values of three biological and three technical replicates. SEM: average of 150 cells. N.a.: not applicable. Significance: * p<0.05, ** p<0.01, *** p<0.001.

**Figure 5.**
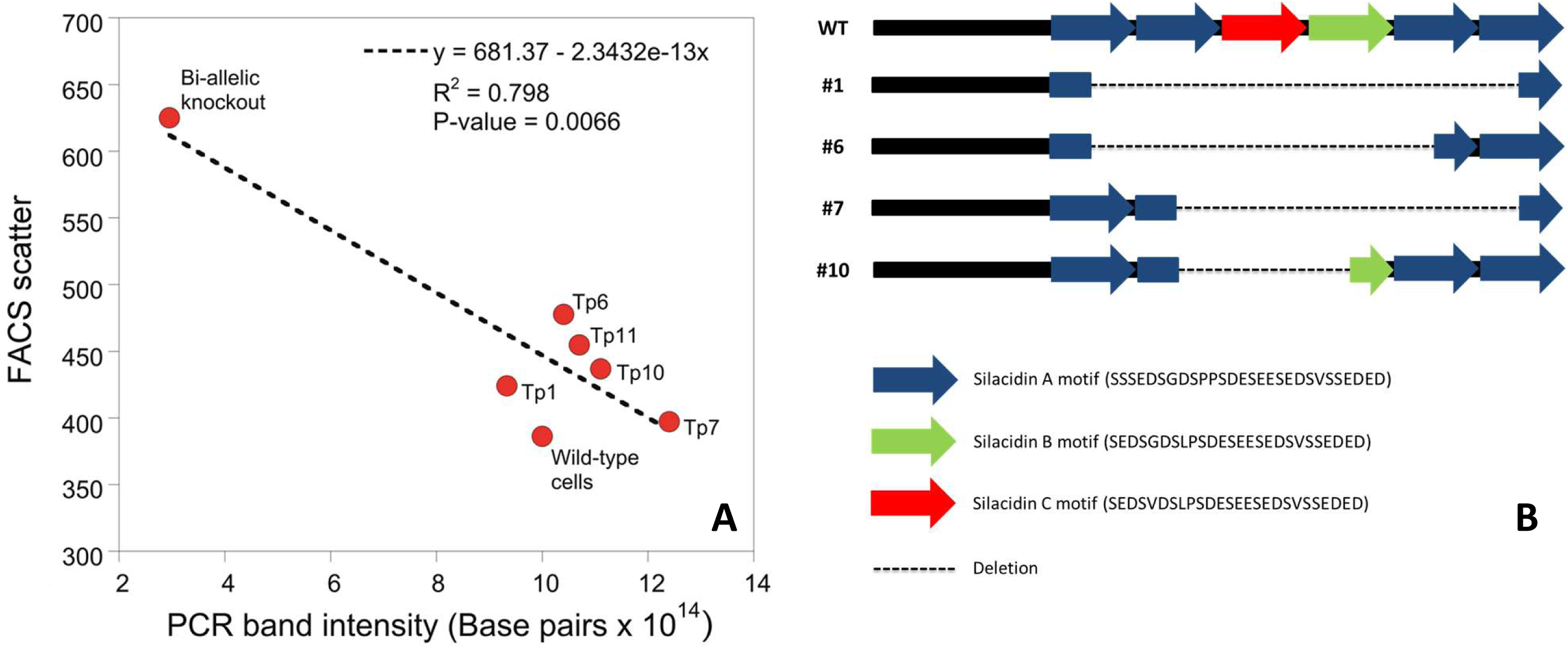
Linking genotype to phenotype in transformant clones. A: Correlation between cell size (measured by FACS) and base pairs per silacidin amplicon (quantitation of PCR bands in Fig. 3C). B: Schematic view of predicted silacidin proteins derived from translating sequencing data in WT and transformant HR clones 1, 6, 7 and 10.

The difference in cell size between WT and HR clones was caused by differences in the diameter of the valves, which are overlapping sections of the cell walls also known as thecae (Fig. 6).

**Figure 6.**
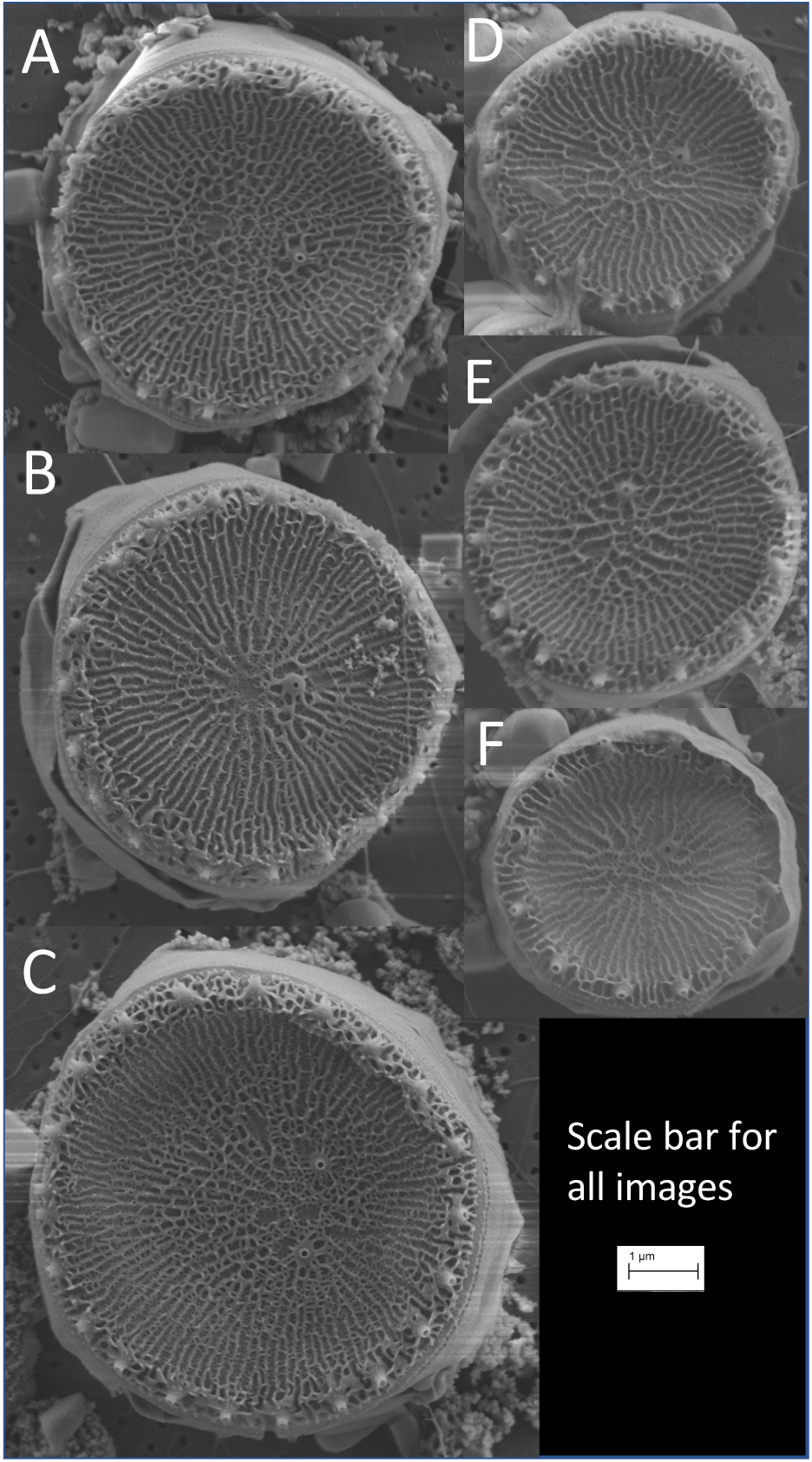
Representative scanning electron microscope images of *T. pseuodonana* valves. Individual cells from HR clone 12 (A-C) and wild-type cells (D-F).

## Discussion

Although the applicability of CRISPR/Cas9 for genome editing in *T. pseudonana* has recently been demonstrated (Hopes et al. 2016), the current work provides the first evidence for highly efficient CRISPR/Cas9-mediated HR in *T. pseudonana*, which is unprecedented for any photosynthetic organism. This surprising result was achieved by designing an exogenous donor matrix to substitute the cell-wall gene silacidin with a resistance cassette encoding the *NAT* gene using Golden Gate cloning. Similar experiments with the diploid diatom *P. tricornutum*, which, however, were based on TALEN-mediated HR, resulted in a maximum HR efficiency of only 27% when taking into account the number of colonies which tested positive for HR on selective plates (Daboussi et al. 2014) (Table 3). In the present study, however, CRISPR/Cas9-mediated HR was detected in 85% of obtained colonies, making it the most efficient incidence of HR reported for any photosynthetic eukaryote with a predominantly diploid life cycle phase. Other photosynthetic eukaryotes with a similar HR efficiency were either haploids such as *Nannochloropsis sp*. or at least had a predominantly haploid life phase such as *Physcomitrella patens*. However, HR in both species was not mediated by nucleases. The highest HR efficiency mediated by any nucleases among all photosynthetic eukaryotes was observed in *P. tricornutum* to date. Thus, the results of our study represent an unexpectedly high rate of HR events overall and compared to NHEJ in particular. Usually, NHEJ is expected to be the primary repair mechanism if Cas9, or any other exogenous nuclease, induces breaks within the target DNA sequence. NHEJ was reported to be preferred over HR in human cells and diatoms (Mao et al. 2008; Daboussi et al. 2014). However, in our study, no evidence for NHEJ was detected despite the use of a sgRNA targeting two sites. Potentially, repair events of a single cut remained undetected by our screening approach but were still responsible for efficient HR (Figs. 3A and B). Based on our sequencing results, the truncated forms of the silacidin gene seemed to be caused by HR between the direct repeats of the silacidin gene leading to ‘looping out’ and deletion of one or more of the six repeats.

**Table 3.**
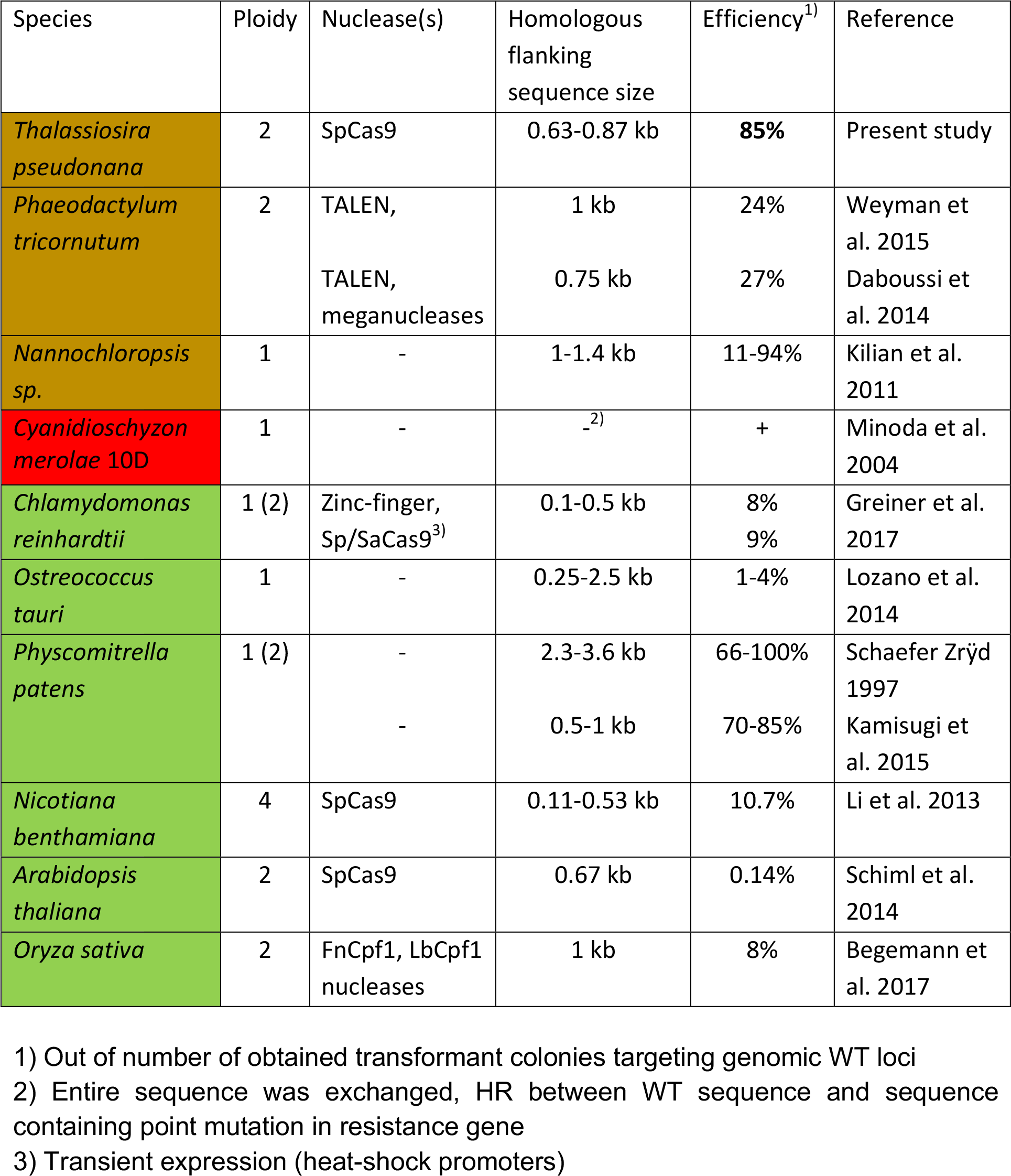
Overview of HR occurrence and efficiency in photosynthetic eukaryotes. Sa: *Staphylococcus aureus*, Sp: *Streptococcus pyogenes*, Fn: *Fransicella novicida*, Lb: *Lachnospiraceae bacterium*. Heterokonta: brown, Rhodophyta: red, Plantae: green.

Interestingly, phenotyping the WT, complete bi-allelic HR, and HR colonies with different truncated versions of the silacidin gene, revealed a significant negative correlation (p-value = 0.007) between the level of mutations caused by HR (genotype) and the cell size expressed as the intensity of light scattering (phenotype) (Fig. 5A). This correlation suggests that the truncated versions of the silacidin gene (shorter bands in Fig. 3C, lanes 1, 6, 7 and 10) were still functional in the HR cell lines. The largest cell size (Table 2) was only observed when the silacidin gene was completely replaced by the FCP:*NAT* cassette as observed for clone 12 (Fig. 3C).

The repetitive nature of the silacidin gene in addition to known post-translational modifications of the encoded protein might explain the significant link between genotype and phenotype. The truncated sequences displayed several different deletions, which nevertheless were still in frame and therefore probably translated into shorter versions of the repetitive WT silacidin protein sequence (Fig. 5B).

Repetitive motifs that undergo extensive post-translational modification such as phosphorylation ((SS)SEDSXDSXPSDESEESEDSVSSEDED) were shown to be the functional units of silacidin proteins responsible for silica precipitation in the cell wall (Fig. 5B;. Even beyond this exact peptide motif, the overall number of serines and acidic amino acids are most likely contributing to the functionality of the truncated versions. These data represent the first example of a direct link between genotype and phenotype with respect to how mutations in a single gene encoding a cell-wall protein impact the cell size of a diatom. Thus, our work will contribute to firmly establishing *T. pseudonana* as a model for bionanotechnology, but also for questions addressing fundamental diatom biology including applications targeted on harnessing the metabolic potential of diatoms.

## Acknowledgements

This work was funded by a grant from the Gordon and Betty Moore Foundation (Grant #4961) in addition to support received from the Natural Environment Research Council (NERC) (NE/K013734/1). AH acknowledges funding from NERC for her PhD studentship.

## Authors’ contributions

TM and AH conceived the project. TM wrote the paper jointly with IG with contributions from NB, AH and AG. NB designed and conducted the laboratory and bioinformatics work with help of IG. LA conducted the cell-size measurements and produced the scanning electron micrographs.

## Figure legends

Figure S1. Alignment of sequencing data of silacidin amplicon in WT, HR clones 1, 6, 7 and 10.

